# Structural variants contribute to pangenome evolution of a plant pathogenic fungus

**DOI:** 10.1101/2021.04.14.439764

**Authors:** Li Guo, Quanbin Dong, Bo Wang, Mengyao Guo, Kai Ye

## Abstract

Genetic variation is the driving force of plant-pathogen co-evolution. Large-scale genetic variations such as structural variations (SVs) often alter genome stability and organismal fitness. However, the pangenomic landscape and functional implications of SVs remain largely unexplored in plant pathogens. Here, we characterized the pangenomic and SV landscape in wheat head blight fungus *Fusarium graminearum* by producing and comparing chromosome-level (average contig N50 of 8.9 Mb) genome assemblies of 98 accessions using a reference-guided approach. Accounting for 29.05% and 19.01% of *F. graminearum* pangenome, respectively, accessory and private genomes are enriched with functions related to membrane trafficking, metabolism of fatty acids and tryptophans, with the private also enriched with putative effectors. Furthermore, using chromosome-level assemblies, we detected 52,420 SVs, 69.51% of which are inaccessible using read-mapping based approach. Over a half (55.65%) of 52,645 merged SVs affected 1,660 protein-coding genes, the most variable of which are involved in fungal virulence, cellular contact and communications. Interestingly, highly variable effectors and secondary metabolic enzymes are co-localized with SVs at subtelomeric and centromeric regions. Collectively, this landmark study shows the prevalence and functional relevance of SVs in *F. graminearum*, providing a valuable resource for future pangenomic studies in this cosmopolitan pathogen of cereal crops.

## INTRODUCTION

Fungal pathogens contribute to a substantial fraction of crop diseases and challenge global food safety, economic and social stability (Savary & Willocquet, 2020). For example, rice blast disease caused by *Magnaporthe oryzae* threatens rice productions worldwide (Dean et al., 2012). Fusarium head blight caused by *Fusarium graminearum* is a devastating disease of wheat and barley causing huge yield and economic losse (Goswami & Kistler, 2004). FHB also threatens human and animal health through mycotoxins such as trichothecenes and the estrogenic zearalenone (Chanda et al., 2016). A major obstacle of battling against many devastating crop diseases including FHB is the constant and rapid evolution of pathogen virulence and drug resistance through gene mutation and natural selection, an inevitable problem further deteriorated by fungicide abuses and resistant cultivar monoculture widely adopted in modern agriculture. Drug resistance in agricultural pathogens also poses dangers to human health through opportunistic fungal infections in immunocompromised individuals (Benitez & Carver, 2019). It is thus necessary to investigate the landscape and function of genetic mutations leading to evolution of fungal traits such as virulence and antifungal resistance, so that effective and environment-friendly strategies can be developed for plant disease prevention and management.

Genetic variants arisen from DNA mutations are the driving force behind evolution (Kronenberg *et al*., 2018) including host-pathogen co-evolution with a boom-and-bust cycle (De la Concepcion *et al*., 2018). Genetic variations come in various forms including single nucleotide polymorphisms (SNPs), small (<50bp) deletions or insertions (Indels) and structural variations (SVs) (>50bp) (Mahmoud *et al*., 2019). Generally, genetic variants modify gene coding or non-coding sequences leading to altered gene functions and ultimately organismal fitness (Kronenberg *et al*., 2018). Because these variants contribute to the formation of genetically diverse populations, any reference genome assembly of a single individual hardly represents the complete genetic information of any species known as pangenome (Parfrey *et al*., 2008). Pangenome represents a non-redundant complement of genome sequences for all individuals within a species (Tettelin *et al*., 2005). First defined in bacteria, pangenome has been conceptually recognized and explored across all major kingdoms ranging from human (Li *et al*., 2010), animals (Li *et al*., 2017; Tian *et al*., 2020), plants (Bayer *et al*., 2020) to bacteria (Ding *et al*., 2018) and fungi such as *Saccharomyces cerevisiae, Candida albicans, Cryptococcus neoformans, Aspergillus fumigatus* (Golicz *et al*., 2016; Peter *et al*., 2018), and also several plant pathogens including *Parastagonospora* spp, *Zymoseptoria tritici* (Plissonneau *et al*., 2018; Syme *et al*., 2018). Therefore, characterization of genetic variants is vital to mapping the pangenomes and understanding the mechanisms of species evolution.

Despite the importance of both small and large variants, our current understanding of fungal genetic variations generally focuses on SNPs that are widely used in population genetics and genome-wide association studies to link genotypes with phenotypes. So far, *F. graminearum* population genetic studies have emphasized on analysis of SNPs. For example, a link between local polymorphisms and pathogen specificity has been identified in *F. graminearum* genome (Cuomo *et al*., 2007). Firasl *et al*. associated SNP diversity with genes crucial for *F. graminearum* phenotype including trichothecene chemotypes and virulence (Talas *et al*., 2012). By contrast, there is to date a lack of both interest and effort in studying large variants such as indels and SVs in population genetic studies of fungal pathogens. However, compared to SNPs, SVs are more likely to disrupt the genome stability and function such as altering gene structure, copy numbers, and gene regulation given their large size (Alonge *et al*., 2020). For example, SVs have already been implicated in development of various genetic disorders in certain human pedigrees or populations (Friedman *et al*., 1994; Nattestad *et al*., 2018). Therefore, a lack of population-wide mapping of structural variants in plant pathogens has led to an underestimation of their genetic diversities as well as impact on fungal pangenome evolution.

The overall lack of SV knowledge in plant pathogenic fungi is largely down to the technical challenges to detect SVs based on widely-used next-generation sequencing (NGS) data due to its small read-length (Mahmoud *et al*., 2019). Although variant detection tools such as *Pindel* (Ye *et al*., 2009), *Delly* (Rausch *et al*., 2012), *Lumpy* (Layer *et al*., 2014) are available, application of these tools to NGS data are mostly ideal for detecting small variants with limited power in large variant discovery. Third-generation sequencing technology (*i*.*e*., Pacific Bioscience or Oxford Nanopore), able to span most repetitive and complex regions in genome assembly and variant detection given the long reads (Mahmoud *et al*., 2019), presents an ideal alternative to identify SVs. However, long-read sequencing remains expensive for variant detection in large-scale population genomic studies of plants and fungi. Recently, an alternative strategy has been proposed for variant detection based on a chromosome-scale reference genome and population-scale resequencing datasets. It involves reference-guided scaffolding of draft genome assemblies from NGS data, followed by assembly-based detection of variants. Several computational tools have been developed for this task including *Ragout2* (Kolmogorov *et al*., 2014) and *RaGOO* (Alonge *et al*., 2019) etc., providing a fast and affordable option to characterize pan-SV landscape at population level. The chromosome-scale genome assemblies also facilitate the analysis of pangenomes for the species being studied.

In this study, we sought to identify SVs in a large collection of *F. graminearum* accessions using chromosome-level genome assemblies, generated by referenced-guided genome scaffolding of NGS-based assembly, followed by SV identification. We also constructed the pangenome of *F. graminearum* based on these assemblies, revealing the contribution of accessory and private genomes to species adaptation. Intersecting the SVs with pangenome components highlighted the important role of SV in the genome evolution and pathogenesis of *F. graminearum*. This study not only presents a valuable resource for future population genomic and pangenomic investigation in this cosmopolitan fungal pathogen, but also demonstrates how SVs could be analyzed in fungal population genomic datasets solely based on NGS.

## MATERIALS AND METHODS

### Sequencing data and quality control

NGS (Illumina paired end) raw data of 104 *F. graminearum* isolates from five countries (China, USA, United Kingdom, France and Australia) around the globe were downloaded from National Center of Biological Information (NCBI) Sequence Read Archives (SRA) (Table S1). The SRA data were then converted to FASTQ format using SRA Toolkit (https://github.com/ncbi/sra-tools). The quality of the FASTQ data were assessed from two perspectives. Firstly, *FASTP* (Chen *et al*., 2018) was used to check the read quality such as base quality, guanine-cytosine (GC) content, adapters etc. of the fastq files, followed by filtering reads with the poor quality and adapters with default parameters settings. Secondly, the software *Sourmash* (Ondov *et al*., 2016) was used to check k-mer distributions of each dataset, finding and filtering out samples with abnormal k-mer frequencies. In total, 98 of 104 samples passed the quality control and these cleaned data were used for the downstream analysis.

### Chromosome-level genome assembly

*SPAdes* (Prjibelski *et al*., 2020) was used to *de novo* assemble the cleaned reads, with the parameters: -k 33,55,77 --careful -t 28, and then the contigs.fasta and scaffolds.fasta were generated. *RaGOO* (Alonge *et al*., 2019) was used to assemble contigs on the chromosome level based on the results of *SPAdes* (Prjibelski *et al*., 2020). The running parameter was -b -t 4-g 100-s-i 0.2, and the fasta file at the chromosome level was obtained. To evaluated the genome assemblies, we run *QUAST* (Gurevich *et al*., 2013) with default parameters.

### Genome annotations and effector prediction

For *F. graminearum* genome annotation, *de novo* gene structure was predicted by *GeneMark-ES* with parameters ‘--ES --fungus’ (Lomsadze *et al*., 2005; Ter-Hovhannisyan *et al*., 2008). A Fusarium gene model was then used to train *AUGUSTUS* v. 3.1 (Stanke *et al*., 2008). *MAKER2* pipeline (Min *et al*., 2017) with *RepeatMasker* v. 4.0.7 (Saha *et al*., 2008) option on to find and mask repetitive elements, was used to find protein-coding genes integrating gene models predicted from *GeneMark-ES* and *AUGUSTUS*, and protein sequences of the *F. graminearum*. The *F. graminearum* putative effectors were predicted as follows: candidate secreted proteins have a secretion signal as determined by *EffectorP* (Sperschneider *et al*., 2018) and have no transmembrane domain as determined by *TMHMM* 2.0 (Krogh *et al*., 2001). Eventually, *WoLF-PSort* v. 0.2 (Horton *et al*., 2007) software was used to estimate the located sites and only those proteins that were credibly positioned in the extracellular space (i.e., extracellular score >15) were included into in the final secretome (Kaundal *et al*., 2010). Small secreted proteins (SSPs) are defined here as proteins that are smaller than 200 amino acids and labeled as ‘cysteine rich’ when the percentage of cysteine residues in the protein was at least twice as high as the average percentage of cysteine residues in all predicted proteins of that organism.

### Variant detection

Structural variant detection was conducted using two different approaches: mapping based approach (MBA) and assembly-based approach (ABA). For MBA, we first mapped NGS short reads to *F. graminearum* PH1 genome using *BWA-mem* (Li & Durbin, 2009), and performed structural variant detection using three mainstream SV callers *Lumpy* (Layer *et al*., 2014), *Delly* (Rausch *et al*., 2012) and *Manta* (Chen *et al*., 2016), followed by merging the detected SV of each caller (only considering SVs that are detected by at least two of four SV callers) using *SURVIVOR* (Jeffares *et al*., 2017). Alternatively, with ABA we aligned each of the 98 chromosome scale genome assemblies against *F. graminearum* PH1 genome, followed by structural variant detection using *Assemblytics* (Nattestad & Schatz, 2016). The chromosome-level genome assembly for each of 98 *F. graminearum* isolates was aligned to the reference genome PH1 using *minimap2* (Li, 2018) with the parameter settings: *minimap2 -k19 -w19 reference*.*fasta contigs*.*fasta*, where “ reference.fasta” and “ contigs.fasta” represents the PH1 reference genome and genome assembly results given by *RaGOO*, respectively. The alignments (.pav files) were then converted to delta format, and then used as input to *Assemblytics* for structural variant discovery with the parameter settings: *assemblytics contigs*.*delta contig_SV 1000000 1 1000000*. Structural variants detected recorded in .bed files as the output of *Assemblytics* were converted to VCF (variant call format) files using *SURVIVOR* v2.0.1. Structural variants of multiple isolates were filtered, compared and merged using *SURVIVOR* to identify common and distinct variants. SNP and indels were identified using Genome analysis tool kit (*GATK*) (DePristo *et al*., 2011). *dN*/*dS* (the ratio of non-synonymous to synonymous substitutions) data were obtained from a previous report by Sperschneider *et al* (Sperschneider *et al*., 2015).

### Structural variant effect analysis

The effects of structural variants on genome functions were analyzed using *ANNOVAR* (Wang et al., 2010). Genome annotation files (.gtf) and VCF files storing structural variant calls and genome coordinates were used as input to *ANNOVAR* for calculating the effects of each structural variant including overlaps with gene coding regions (introns and exons), UTRs, intergenic regions etc. The fungal genes affected by structural variants were obtained by overlapping the gene annotation information with the variant information stored in .bed files given by *Assemblytics* using *Bedtools* (Quinlan & Hall, 2010). A threshold of 10% or more in gene coding regions overlapping with any structural variant was used to identify genes affected by the variant.

### Pangenomic analysis

The pangenome analysis was conducted using two different approaches: genome-based and gene-based approach. For genome-based approach, ppsPCP pipeline (Tahir Ul Qamar *et al*., 2019) was used for pan-genome analysis to find full complement of genome sequences from all 98 genomes with default parameters. For the gene-based approach, we used protein sequences of all 98 isolates and PH1 to identify ortholog groups (orthogroups) shared by all proteomes, among different proteomes and unique to each proteome using *OrthoFinder* (Emms & Kelly, 2019). Core genome is defined as orthogroups present in all isolates, whereas accessory genome is defined as orthogroups shared by some but not all isolates. Private genome is defined as orthogroups unique to each isolate. The three parts of pangenomes were compared with genes encoding for effectors, carbohydrate-degrading enzymes, virulence factors (PHI-base records (Urban *et al*., 2020)) and trichothecene biosynthetic enzymes to evaluate the evolution of these gene functions in *F. graminearum*. The pangenome components were also compared with genes affected by structural variants to assess the contribution of the variants to these gene functions and fungal evolution. Pangenome openness was determined by fitting the pangenome profile curve model: *y= AxB + C* (Tettelin *et al*., 2005), where y and x represent pangenome size and genome number respectively, and A, B and C are filting parameters.

### Functional enrichment analysis

For gene function annotation, KEGG pathway analysis was performed using KOBAS3.0 (Xie *et al*., 2011), protein domain was annotated by InterProScan (Jones *et al*., 2014), and Gene Ontology was annotated by BLAST2GO (https://www.blast2go.com/), and then enrichment analysis was completed by TBtools (Chen *et al*., 2020).

### Data availability

The genome assemblies and variants reported in this paper have been deposited in the Genome Sequence Archive in National Genomics Data Center, China National Center for Bioinformation / Beijing Institute of Genomics, Chinese Academy of Sciences, under the BioProject ID PRJCA004286 and accession numbers WGS018715-WGS018812 that are publicly accessible at https://bigd.big.ac.cn/gsa.

## RESULTS

### Chromosome-level genome assembly of 98 *F. graminearum* accessions

To reconstruct the pangenome and identify structural variants in *F. graminearum*, we produced chromosome-level genome assemblies for a collection of *F. graminearum* accessions via a reference-guided approach. We first downloaded from public domains NGS data for 104 *F. graminearum* isolates originally sampled from five countries: China (CN), the United States (US), France (FR), United Kingdom (UK) and Australia (AUS) (Table S1). Quality of these NGS data were assessed, followed by removing six problematic datasets (showing abnormal Kmer frequencies) and poor-quality reads and sequence adapters, yielding a total of 98 high quality datasets including 60, 24, 6, 4 and 4 from US, AUS, FR, UK and CN, respectively (Figure 1A). Cleaned reads were then *de novo* assembled by *SPAdes* to generate 98 draft genome assemblies (Figure 1B; Table S1) with genome sizes ranged from 34.3Mb to 37.4Mb and an average GC content of 48.20%. Unsuprisingly, these assemblies were overall fragmented with the number of contigs ranging from 72 to 805 (Figure 1B and 1C), and contig N50 ranging from 93.8kb to 2.3Mb (Figure 1C).

**Figure 1.**
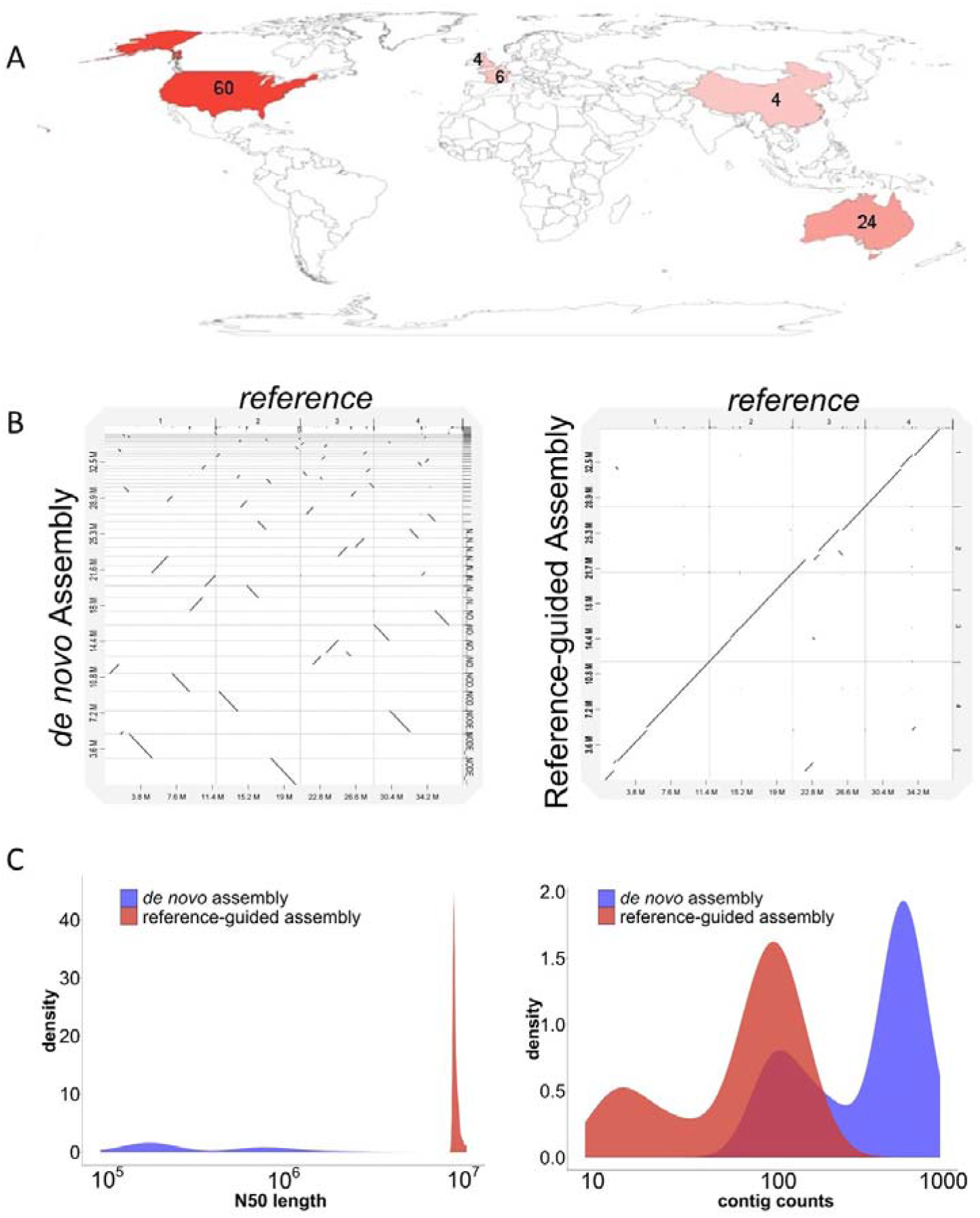
Geographic distribution and genome assembly of 98 *Fusarium graminearum* accessions. **A**. World map displaying the countries of origin for the *F. graminearum* accessions included in this study. The color scale is proportional to the number of accessions marked on the map. **B**. Whole genome alignments of *F. graminearum* reference genome PH1 against the genome assembly using Illumina short reads alone (left) and using *RaGOO* to perform a scaffolding based on the NGS assembly (right), using UK2999 isolate as an example. **C**. Density distribution of contig counts (left) and contig N50 (right) for the 98 genome assemblies using short reads alone (blue) or using reference-guided assembly of the short reads (red).

High-quality genome assemblies are needed for optimal pangenome construction and efficient SV identification based on whole genome alignments. Recently, several algorithms such as *RaGOO* (Alonge *et al*., 2019) for reference-guided genome assembly have been developed to scaffold contig-level assemblies into chromosome-level assemblies using a reference genome. From the NGS-based draft genomes of the 98 isolates, we further generated chromosome-scale genome assembly for each isolate using *F. graminearum* PH1 reference genome as a guide (Figure 1B). We obtained 98 final genome assemblies of high contiguity with contig N50 ranging from 8.3Mb to 10Mb (Figure 1C), a significant leap of quality over the draft assemblies (Table S1). We also observed that the draft contigs of each isolate could not be fully aligned into four chromosomes of PH1 genome, suggesting that each isolate has underwent substantial evolution carrying unique genome sequences. It demonstrated the need to characterize the fungal pangenome because any individual genome is insufficient to represent the genetic information in the whole species.

### Pangenomic analysis of *F. graminearum*

We recovered the *F. graminearum* pangenome sequence by a genome-based approach from the 98 chromosome-scale genome assemblies using ppsPCP pipeline (Tahir Ul Qamar *et al*., 2019). First, each genome assembly was iteratively compared with the PH1 reference genome, followed by the presence-absence variation identification via scanning unique sequences (>100bp) of each accession relative to the reference genome. For each iteration, the unique sequences and the reference genome were merged into a non-redundant sequence file. The process was repeated for all 98 accessions to complete the pangenome construction for *F. graminearum*. The final pangenome size of the 98 accessions is 42.6Mb, about 5.6Mb larger than the PH1 reference genome. These extra sequences encoded a total of 1,203 protein-coding genes, and functional enrichment showed that they were mostly significantly enriched in pathways such as carbohydrate, fatty acid and tyrosine metabolism, transporters (Figure S1). Fatty acid, carbohydrate and amino acid metabolism produces primary metabolites that are not only essential for fungal cellular functions, but also precursors for fungal secondary metabolism (Chroumpi *et al*., 2020).

For any species, pangenome typically consists of gene sets conserved in all, some or none of the isolates, which are defined as core, accessory and private genomes, respectively. To systematically identify the core, accessory and private genomes in *F. graminearum* pangenome, we first predicted the protein-coding genes from the chromosome-scale genomes of the 98 *F. graminearum* accessions using *AUGUSTUS* (Stanke *et al*., 2008) based on Fusarium-specific gene model (Table S1). *Orthofinder* was then used to identify orthologs between PH1 and 98 samples, classifying genes into 15,408 orthogroups, among which 8,003 (51.94%) were present in all samples defined as the core genomes. Additionally, 2,928 (19.01%) orthogroups associated with a single accession, defined as private genomes. Finally, the remaining 4,476 (29.05%) orthogroups associated with at least two but not all accessions were defined as accessory genomes (Figure 2A and 2C). We found that the pangenome size increased before reaching a plateau as the number of accessions increased, but the size of core genomes decreased (Figure 2B), suggesting that *F. graminearum* has a closed pangenome. Interestingly, we found significant smaller dN/dS ratios were associated with the *F. graminearum* core genes than with the accessory genes and private genes, suggesting a different selection pressure likely being exerted on the three types of genomes (Figure 2D). Furthermore, functional enrichment showed that the accessory genes were enriched in membrane trafficking (SNARE mediated vesicle trafficking, exocytosis and autophagy), ribosome and protein translation. Private genes were enriched in transcription factors, metabolism of amino acids (valine, leucine and tryptophan) and fatty acids (Figure 2E), consistent with the finding using genome-based approach (Figure S1). By contrast, core genomes were enriched in pathways related to the basic metabolism and house-keeping cellular processes (Figure 2E). Collectively, the pangenomic analysis indicated that *F. graminearum* field populations have evolved accessory and private genomes with stronger diversifying selection compared to core genomes, reflecting the pangenome evolution behind the fungal adaptation.

**Figure 2.**
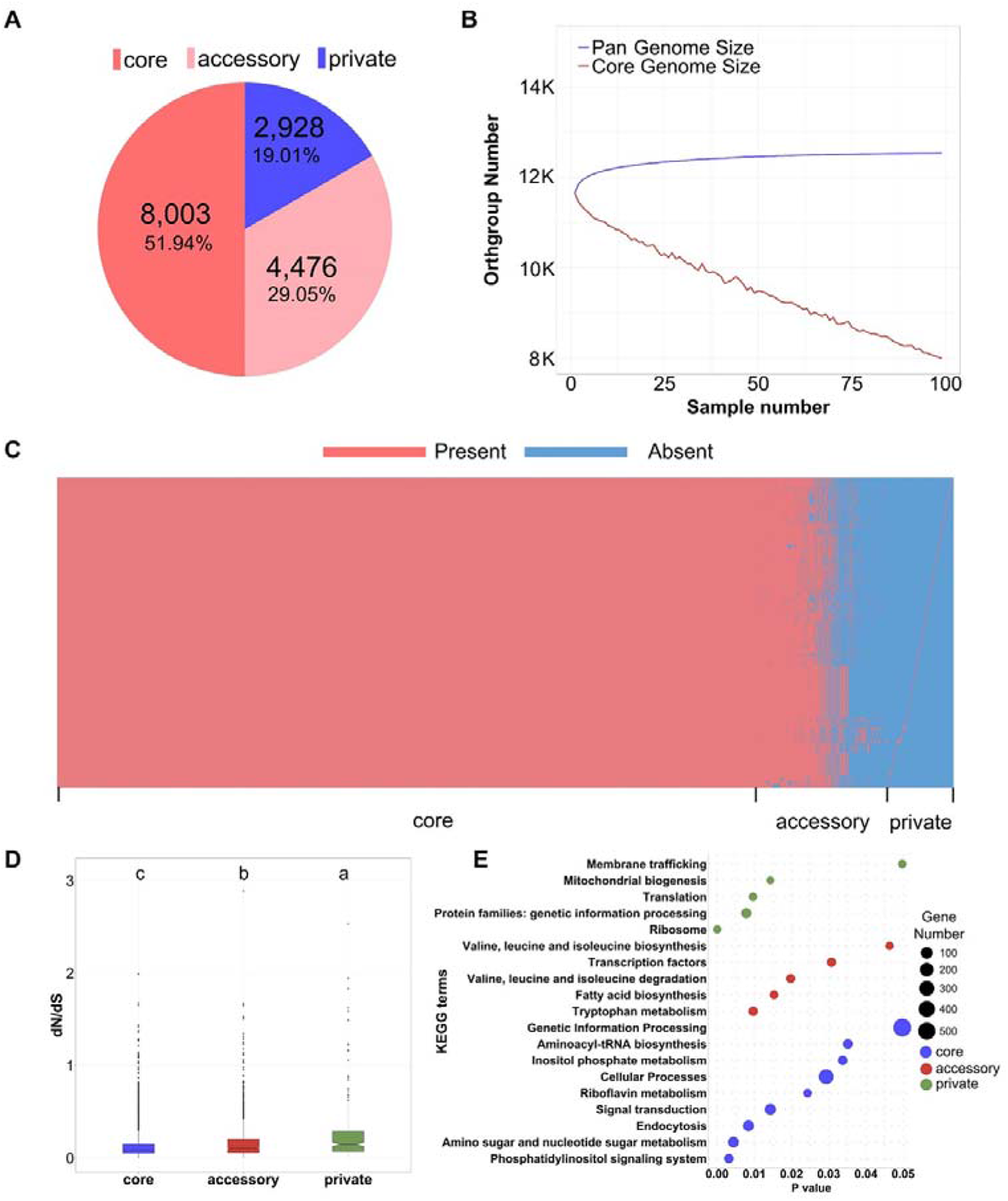
Pan-genome analysis of *Fusarium graminearum*. **A**. Core, accessory and private genomes represent 51.94%, 29.05% and 19.01% of *F. graminearum* pan-genome, respectively. **B**. Variation of gene families in the pan-genome and core-genome along with an additional *F. graminearum* genome. **C**. The number of genes counted for each pan-genome composition (core, accessory and private) in 98 individual genomes. **D**. Boxplot of *dN/dS* ratio (nonsynonymous substitution rate divided by synonymous substitution rate) distribution for *F. graminearum* genes located on each pan-genome composition (core, accessory and private). The lower-case letter a, b and c represents the significant difference (p < 0.05) using Student’s t-test. **E**. A bubble plot summarizing the functional enrichment analysis of each composition of *F. graminearum* pangenome. Y-axis and X-axis denotes the enriched KEGG terms and p value (p <0.05).

### Mapping structural variants in *F. graminearum*

Genetic variants play a central role in genome evolution. With the identified *F. graminearum* pangenome, we are curious about what genomic variations each accession went through to shape the current fungal genome. We characterized the structural variations (SVs) in all 98 *F. graminarum* accessions, as SNPs and indels have already been reported in these isolates previously by others (Cuomo *et al*., 2007; Talas *et al*., 2012). More importantly, SVs are genetic variations typically larger than 50bp such as deletions, insertions, inversions, and translocations, and tend to have more severe consequences to genome stability and organismal fitness (Medvedev *et al*., 2009; Escaramís *et al*., 2015). Here, we focused on detecting large deletions and insertions, two most common SV types, in 98 *F. graminearum* isolates using two different approaches: mapping based approach (MBA) and assembly-based approach (ABA). For MBA, we first mapped NGS short reads to *F. graminearum* PH1 genome using *BWA-mem* (Li & Durbin, 2009), and performed structural variant detection using three mainstream SV callers *Lumpy* (Layer *et al*., 2014), *Delly* (Rausch *et al*., 2012) and *Manta* (Chen *et al*., 2016), followed by merging variants (only considering SVs that are detected by at least two of three SV callers) using *SURVIVOR* (Jeffares *et al*., 2017). Alternatively, for ABA we aligned each of the 98 chromosome-scale genome assemblies against *F. graminearum* PH1 genome, followed by structural variant detection using *Assemblytics* (Nattestad & Schatz, 2016).

In total, the MBA method detected 10,253 SVs (> 50bp) including 10,118 deletions and 135 insertions from 98 *F. graminearum* isolates (Figure 3A). Conversely, the ABA method discovered a total of 52,420 SVs including 30,191 insertions (57.59%) and 22,229 deletions (42.41%) (Figure 3A). The fact that more SVs were detected by ABA than by MBA showed the power of chromosome-scale genome assemblies used for SV discovery. A comparison of SVs found that 8,855 SVs were captured by both MBA and ABA, occupying 86.36% and 16.89% of total SVs discovered by MBA and ABA, respectively (Figure 3A). Interestingly, 69.51% SVs (57.15% deletions, 99.55% insertions) detected by ABA were not detected by MBA. The size distribution showed that smaller and larger SVs are more detectable by MBA and ABA, respectively (Figure 3B). Harnessing the strength of both methods, we obtained a merged SV callset by incorporating variants identified by MBA and ABA, yielding a total of 52,645 SVs (Figure S2) for *F. graminearum*, including 23,614 deletions and 29,031 insertions which were used for downstream characterization of their population landscape and functional effects. Interestingly, SVs tend to be clustered at subteleomeric and centromeric regions of *F. graminearum* genome, although SVs were distributed throughout the genome (Figure 3C), consistent with previous reports that SVs occur more frequently in highly complex genomic regions (Sudmant *et al*., 2015).

**Figure 3.**
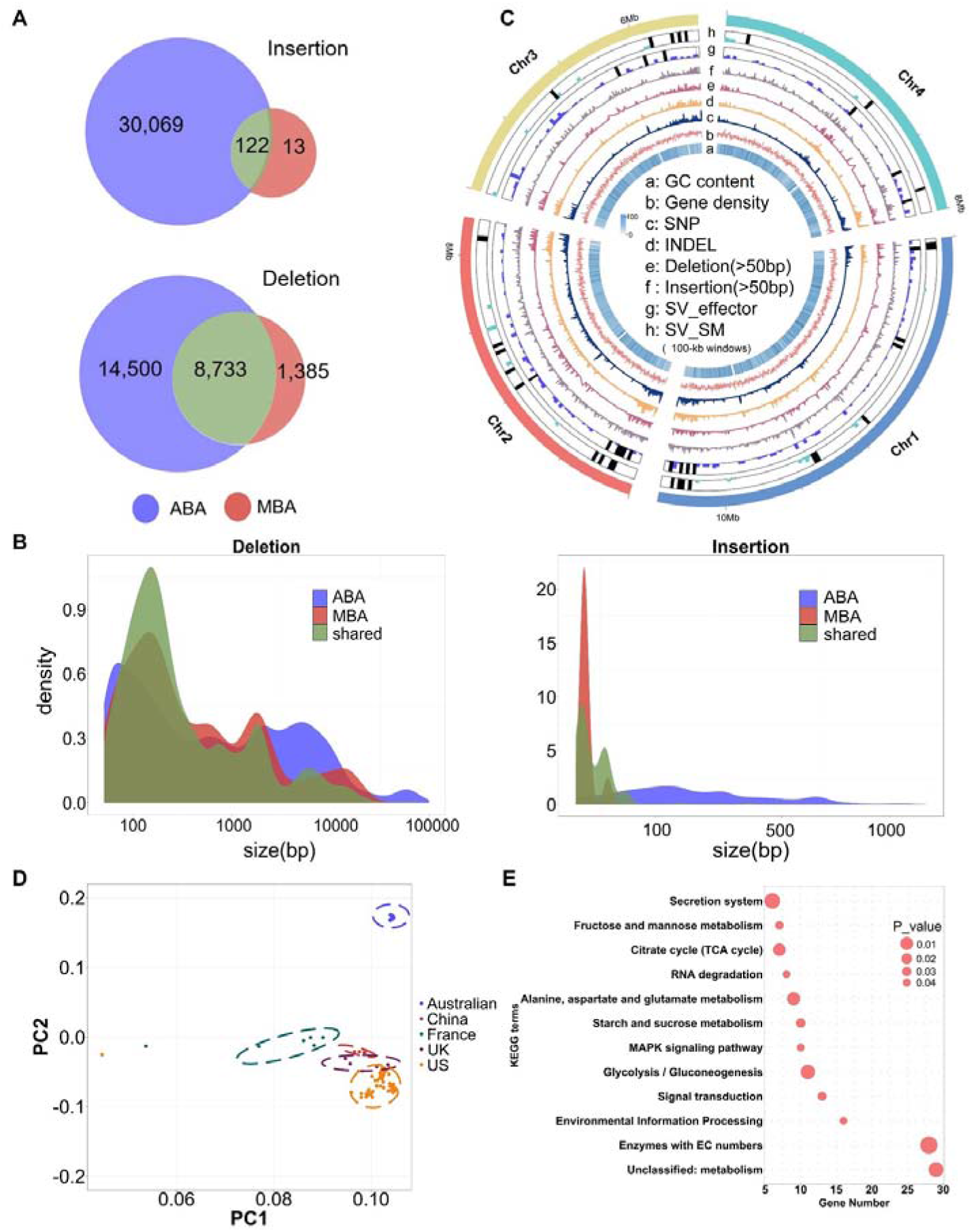
An overview of structural variant landscape in 98 *Fusarium graminearum* accessions. **A**. Comparison of *F. graminearum* structural variants detected using two different approaches: mapping-based approach (MBA) and assembly-based approach (ABA). **B**. The size distribution of structural variants showed that smaller and larger structural variants are more easily detectable by MBA and ABA, respectively. **C**. Genome circos plot displaying the distributions of key genomic features for *F. graminearum*. (a-h) GC content, Gene density, SNP density, indel density, structural variant (SV-deletion, SV-insertion) density, effector and secondary metabolic (SM) gene density calculated in 100-kb windows. Black bars (g and h) represent the highly variable effectors and SM genes intersected with structural variants among at least 80% of *F. graminearum* accessions. **D**. Principal components analysis of the structural variants and geographical locations based on a presence/absence matrix of the 98 accessions. **E**. Kyoto Encyclopedia of Genes and Genomes (KEGG) pathway analyses of SV genes.

Biosynthesis of trichothecene mycotoxins is controlled by *Tri* gene cluster in *F. graminearum* and other trichothecene-producing species (Gauthier *et al*., 2015). Three trichothecene chemotypes have been found in natural isolates of *F. graminearum*: 15-acetyl-deoxynivalenol (15ADON), 3-acetyl-deoxynivalenol (3ADON), and nivalenol (NIV). Studies have shown that gene presence and absence variation within the cluster leads to the fungal chemotypic diversity. In current study, we detected a large deletion event (2,379bp) contributing to the loss of *Tri7* gene in all 3ADON and NX2 chemotype, but not 15ADON chemotype of *F. graminearum* accession from USA (Figure S3). This is consistent with current knowledge that *Tri7*, a trichothecene biosynthesis gene encoding an acetylesterase catalyzing a C-4 oxigenation essential for T2-toxin production in *F. sporotrichioides* (Brown *et al*., 2001), is a pseudogene in *F. graminearum* 15ADON chemotypes and absent in 3ADON chemotypes (Rep & Kistler, 2010). However, this deletion event was not observed in *F. graminearum* accessions from China, France, Australia and England. In addition, we also detected a large segment of deletion (7,640bp) contributing to the loss of *Tri4, Tri5* and *Tri6* genes in 16 of 24 accessions from Australia, which are deletion mutants of the three genes generated using CRISPR-cas9 genome editing in *F. graminearum* (Table S2)(Gardiner & Kazan, 2018). The fact that these deletion events are consistent with the previous reports or prior knowledge indicates the reliability of the structural variant detection procedure in this study.

With the merged SVs, we further examined their population distributions and effects on coding genes. First, a principle component analysis using a SV presence/absence matrix revealed that the 98 isolates belonged to distinct clusters that overall correspond to their geographical regions (Figure 3D). Second, we found the UK isolates and US isolates had the lowest and highest number of SVs per sample, respectively (Figure S4), although this discrepancy of SV frequency could well be a result of insufficient sampling of the *F. graminearum* population of UK compared to US regions. Third, the genome-wide distribution of SVs showed that majority (84.4%) of SVs intersected with gene exonic regions and their upstream and downstream regulatory regions (Figure S5). These SVs affected a total of 1,660 protein-coding genes *F. graminearum* enriched with pathways such as signal transduction and energy metabolism (Figure 3E), suggesting potential disruptive effects of SVs on the gene function and potential fitness. Lastly, the number of common SVs between isolates gradually decreased as the number of compared isolates increased. For instance, 1,660, 597, 145 and 8 protein-coding genes (1kb flanking each side) intersected by SVs were shared by at least 2, 10, 50 and 90 isolates (Table S3). We further identified highly variable genes among 98 accessions intersecting with the greatest number of SVs. The top 20 highly variable genes encode proteins involved in cell contact during mating (agglutinin like proteins), cell surface associated proteins (Mucins), myosins and kinesin proteins, virulence-associated proteins and 2OG-Fe oxygenase etc. (Table 1), suggesting these highly variable genes in *F. graminearum* pangenomes are likely associated with virulence, fungal cell communications and interactions with either other cells, or the environment.

**Table 1.** Summary of the 20 most variable genes intersecting with structural variants and their annotated functions in *Fusarium graminearum* pangenome. The number of variants is the total number of structural variants intersecting with the protein-coding sequene and its 1kb flanking region.

| Gene ID | Annotations | Number of variants | Number of accessions with variants |
| --- | --- | --- | --- |
| FGRAMPH1_01G05643 | agglutinin-Like Protein | 568 | 93 |
| FGRAMPH1_01G15613 | agglutinin-Like Protein | 489 | 94 |
| FGRAMPH1_01G27087 | agglutinin-Like Protein | 232 | 85 |
| FGRAMPH1_01G21813 | myosin light chain kinase | 187 | 89 |
| FGRAMPH1_01G25011 | mucin 1, cell surface associated (MUC1) | 185 | 85 |
| FGRAMPH1_01G15427 | ankyrin-3 protein | 184 | 93 |
| FGRAMPH1_01G12267 | mucin 22 protein | 157 | 85 |
| FGRAMPH1_01G25295 | virulence-associated lipoprotein MIA | 122 | 84 |
| FGRAMPH1_01G13139 | vacuolar carboxypeptidase | 118 | 83 |
| FGRAMPH1_01G22029 | nucleoside phosphorylase | 117 | 81 |
| FGRAMPH1_01G08911 | Extracellular serine/threonine-rich Protein | 116 | 84 |
| FGRAMPH1_01G08231 | cell surface proteins containing the conserved peptide motif (LPXTG) | 113 | 82 |
| FGRAMPH1_01G12003 | kinesin light chain | 109 | 81 |
| FGRAMPH1_01G28273 | peptidase c14 | 108 | 69 |
| FGRAMPH1_01G27923 | 2OG-Fe oxygenase | 104 | 87 |
| FGRAMPH1_01G11565 | Unknown protein | 103 | 73 |
| FGRAMPH1_01G28289 | kinesin light chain | 102 | 96 |
| FGRAMPH1_01G04545 | zinc finger transcription factor | 98 | 93 |
| FGRAMPH1_01G10821 | SNF5-component of SWI SNF transcription activator complex | 97 | 49 |

### Impact of SVs on *F. graminearum* pangenome and pathobiology gene functions

We next investigated how much SVs may have shaped *F. graminearum* pangenomes, by examining the fractions of genes affected by SVs associated with core, accessory and private genomes for each accession. Compared to the proportion of core (52%), accessory (29%) and private (19%) genes in pangenome (Figure 2A), 45%, 29% and 26% genes affected by SVs belong to core, accessory and private genomes, significantly overrepresented on private genomes but underrepresented on core genomes (Figure 4A; Table 2). This suggests a clear skewed contribution of SVs (large deletions and insertions) towards the evolution of private and accessory genomes, compared to core genomes in *F. graminearum*. As such, SVs would have caused extensive gene loss and gain in the fungal populations, leading to a diverse range of dispensable gene content in different accessions. Conversely, the under-representation of SV-affected genes in core genomes might be a consequence of purifying selection against disrupting conserved genes, many of which perform essential house-keeping functions.

**Table 2.** A summary of the core, accessory and private gene fractions in *Fusarium graminearum* pangenome (Global), SV-affected genes (Pan-SV), and genes belonging to four different functional groups (effectors, CAZyme, SMG and TF). Underneath the fractions are p-values given by two-tail Fisher’s exact tests conducted to determine the statistical significance of gene enrichment. SV: structural variants. O: overrepresented. U: underrepresented. N: nonsignificant. NA: nonapplicable. SMG: secondary metabolic genes. TF: transcription factors.

| Number of genes |  | Global | Effector | CAZYme | SMG | TF |
| --- | --- | --- | --- | --- | --- | --- |
| Pangenome<br>(15,407) | Core<br>(8,003) | 51.94% | 32.36%<br>$p = 2.3\text{e-}08$<br>(U) | **68.98%**<br>$p = 1.2\text{e-}06$<br>(O) | 56.03%<br>$p = 0.2549$<br>(N) | **71.01%**<br>$p = 0.0108$<br>(O) |
| | Accessory<br>(4,476) | 29.05% | 22.60%<br>$p = 0.01155$<br>(U) | 30.10%<br>$p = 0.6724$<br>(N) | 33.62%<br>$p = 0.0647$<br>(N) | 28.99%<br>$p = 0.9817$<br>(N) |
| | Private<br>(2,928) | 19.01% | **45.03%**<br>$p < 2.2\text{e-}16$<br>(O) | **0.92%**<br>$p < 2.2\text{e-}16$<br>(U) | 10.34%<br>$p = 1.1\text{e-}06$<br>(U) | 0%<br>NA |
| Pan-SV<br>(1,660) | Core<br>(842) | 50.72%<br>$p = 0.6084$<br>(N) | 26.15%<br>$p = 0.54$ (N) | 61.90%<br>$p = 0.6839$<br>(N) | 53.68%<br>$p = 0.8471$<br>(N) | 65.52%<br>$p = 0.9248$<br>(N) |
| | Accessory<br>(659) | **39.70%**<br>$p = 2.1\text{e-}10$<br>(O) | 32.31%<br>$p = 0.2344$<br>(N) | 38.10%<br>$p = 0.42$ (N) | 37.50%<br>$p = 0.6084$<br>(N) | 34.48%<br>$p = 0.8212$<br>(N) |
| | Private<br>(159) | 9.59%<br>$p = 4.3\text{e-}16$<br>(U) | 41.54%<br>$p = 0.8282$<br>(N) | 0%<br>NA | 8.82%<br>$p = 0.7388$<br>(N) | 0%<br>NA |

**Figure 4.**
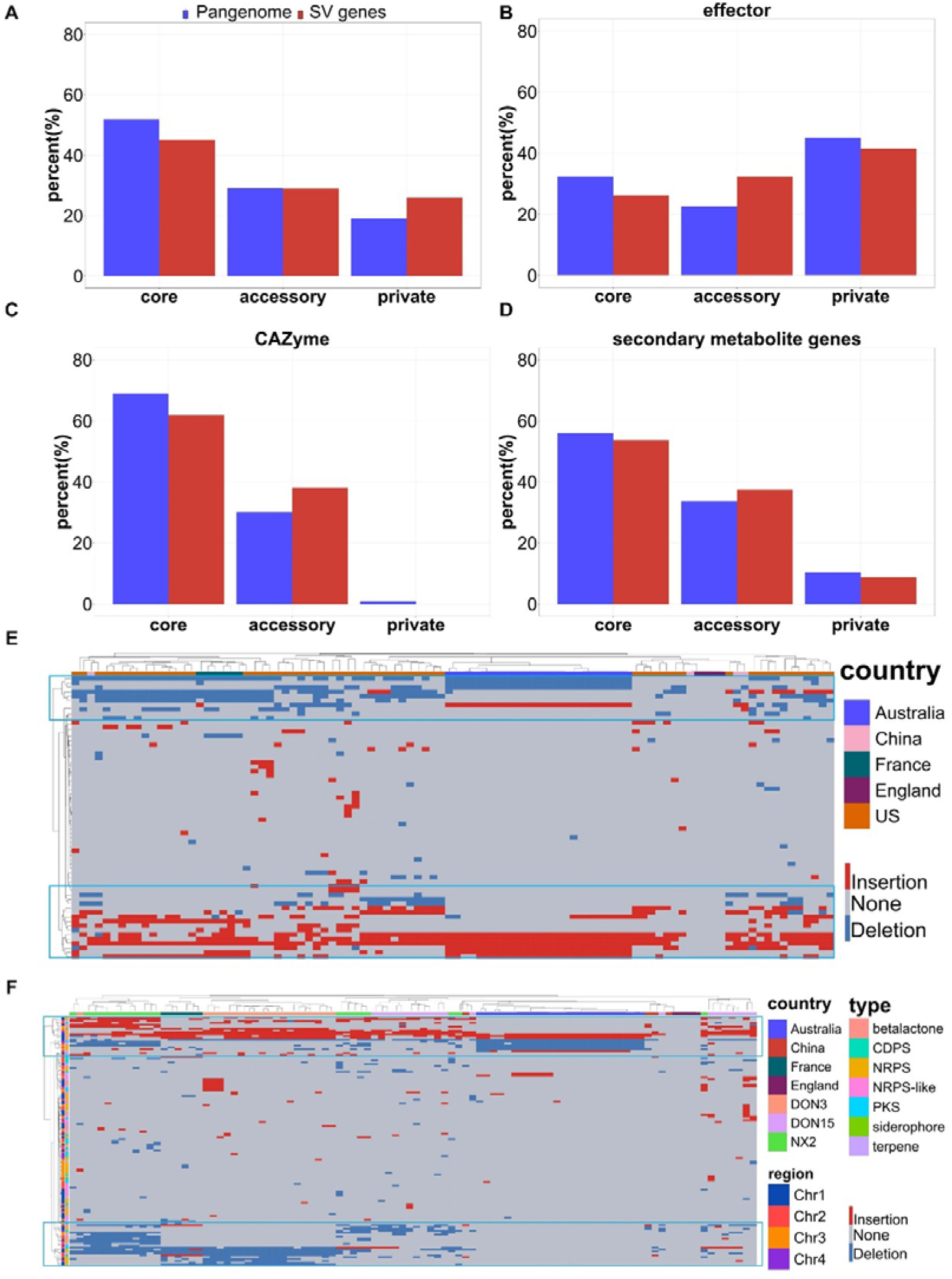
Structural variations contribute to accessory genome evolution in *F. graminearum*. **A**. Proportions of genes affected by structural variants (SV) across the pangenome. **B-D**. Pan-SV categories of carbohydrate-active enzymes (**B**), effectors (**C**) and secondary metabolite (SM) gene clusters (**D**). **E-F**. Heatmaps showing SV frequency of effector (**E**) and secondary metabolic (**F**) genes.

Next, we examined how structural variants have affected specific groups of genes that are associated with pathogenesis or secondary metabolism of *F. graminearum*, including carbohydrate-active enzymes (CAZYme), effectors, secondary metabolic gene clusters and transcription factors. For each of these gene groups, we performed statistical test (Fisher’s exact tests) (Table 2) to determine whether their distribution on each compartment (core, accessory and private) of pangenome significantly deviated from a random distribution of three pangenomic compartments, followed by testing whether such distribution also significantly deviated from the distribution of these gene groups intersecting with SVs on each compartment of pangenome (Table S4). For example, we predicted 584 effector proteins in *F. graminearum* pangenome, small secreted fungal proteins that typically promote pathogenesis, of which 32%, 23% and 45% located in core, accessory and private genome, respectively (Figure 4B), with private genome significantly enriched with effectors (Table 2). We found 65 effectors intersected with SVs, of which 26%, 32% and 42% belong to core, accessory and private genome, respectively (Figure 4B), without enrichment on any compartment (Table 2). Similarly, we analyzed SV impact on 29 transcription factors (TFs), a list of 764 *F. graminearum* CAZYme-encoding genes (Figure 4B) downloaded from dbCAN meta server (Zhang *et al*., 2018), and 696 secondary metabolic genes (SMG) (Figure 4C) we predicted using antismash. The results show that no deviation of distribution was observed for SMGs, global or SV-affected, on any compartment (Table 2). Although TFs and CAZYmes are overall enriched on core genome, no significant enrichment of SV-affected TFs or CAZYmes was found on any compartment (Table 2). Despite such a lack of significant enrichment, an increased proportion on accessory compartment was found for SV-affected SMG (33.62%) and CAZYmes (30.10%) compared to pangenomic ratio (29.05%), suggesting that SVs have contributed to increased variability of these proteins among *F. graminearum* isolates.

Finally, we showed that SMG clusters and effectors harbor substantial structural variations among isolates across different countries (Figure 4E and 4F). We found 22 (33.85%) effectors and 40 (29.41%) SMGs are affected by a deletion or insertion in at least ten isolates, respectively. These highly variable SMGs and effector genes are mostly located at subteleomeric and centromeric regions of chromosomes, consistent with the genomic distribution of SVs (Figure 3C, track g-h). Given likely associations of CAZYmes, effectors and SMG clusters to fungal pathobiology, and the disruptive effects of structural variations on the coding and flanking sequence of these genes, our results indicate that the pathogen pangenome is likely experiencing rapid evolution in these genes allowing the fungus to adapt to host and environmental cues.

## DISCUSSION

The landscape and functional roles of structural variants in fungal pathogens remain an overall uncharted area of research in plant pathogens. Focusing on *F. graminearum*, one of the most researched plant fungal pathogens, we for the first time performed systematic identification of large-scale genome structural variants in a collection of 98 fungal isolates with resequencing data. Knowledge-wise, our study have made new discoveries in three major aspects. Firstly, through reference-guided genome assembly and alignment followed by variant detection, we discover that structural variants are prevalent in *F. graminearum* field populations. Secondly, we show that many of these deletion and insertion variants co-localize with coding genes and thus may disrupt their normal functions. The most highly variable genes (found in over 80% of the *F. grmainearum* accessions analyzed in this study) caused by SVs are involved in agglutin proteins, mucins and kinesins that mediate cell to cell contact and communications during mating or interaction with environment. A high proportion of isolates carrying these mutations indicates pathogen adaptation to surrounding cells or environment is likely under strong selection. Thirdly, although these variants can be found throughout the genome, a high density of SVs is associated with genomic regions near centromere and telomeres. SVs in these highly polymorphic regions intersected with genes encoding putative effectors and secondary metabolic enzymes. Whether SVs play similar roles in evolution of other fungal pathogens of plants and humans would be intriguing to investigate.

Our study also showcased a computational strategy to characterize SVs of plant pathogenic fungi from large populations. The technical challenge of structural variation detection using short reads has been a major reason why these variants are left unnoticed in *F. graminearum*. In this study, we showed that the assembly-based method detected 44,569 structural variants that are inaccessible to traditional read-mapping method, highlighting the limitation of large variant detection based on short reads. Recently, variant callers are being developed to identify SVs in human samples based on single-molecule sequencing data (PacBio and Oxford Nanopore). Therefore, plant and fungal structural variant detections are bound to be improved using these long-read sequencing data given their advantage in detecting large and complex variants, although the cost of producing and analyzing these data from a massive plant or fungal populations remains a tremendous challenge for most large-scale population genomics studies so far. The approach (reference-guided assembly followed by SV detection) we adopted in this study enabled the SV analysis solely based on short reads, proving its efficacy working with population scale resequencing data in pathogenic fungi. With the cost of sequencing continuously plummeting in the near future, it will be possible to obtain long-read-based fungal resequencing data from hundreds or thousands of field isolates or experimental strains to reveal a more complete pangenomic and pan-SV landscape.

*F. graminearum* SVs detected in this work represent a valuable resource for future population genomic and pangenomic studies in this cereal pathogen, which is important for two reasons. First, the prevalence of large scale genome variants in *F. graminearum* genome clearly shows the inadequacy of a single reference genome in population genetic studies, since it tends to introduce geographic bias in interpreting the genomic data. A pangenomic database integrating all types of variants is essential to a more robust interpretation of genetic variations genotyped in various *F. graminearum* populations. Second, failure to characterize the full spectrum of genome variants by missing the structural variants represents a blind spot for discovering the genotype and phenotype associations in *F. graminearum*. Despite the effects of SNPs in gene expression and regulation, they are less disruptive to gene functions and phenotypes than large-scale variations such as SVs and chromosomal aberrations. Therefore, it’s critical to take into consideration the impacts of a broader spectrum of variants for identifying the causal mutations behind trait evolution such as tolerance of antifungal drugs or evasion of host resistance.

In conclusion, we have produced genome assemblies for a large collection of *F. graminearum* isolates, based on which the fungal pangenome and structural variants were comprehensively analyzed. Our study demonstrates that SVs are ubiquitous in *F. graminearum* genomes disrupting functions of genes possibly associated with pathogenesis and secondary metabolism, providing insights into the fungal genome evolution. The computational strategies and structural variant resources developed by this study will be valuable to future population genetic researches of *F. graminearum* and other plant pathogenic fungi.

## AUTHOR CONTRIBUTIONS

LG and KY conceived and designed the project. LG, QBD and MG performed the quality control, variant detection and pangenome analysis. WB conducted the genome assembly and annotation. LG, QBD and KY wrote the manuscript. All authors revised and approved the manuscript.

## ACKNOWLEDGEMENT

This project was supported by the National Natural Science Foundation of China (31701739, 31970317 and 3167372) and National Key R&D Program of China (2017YFC0907500, 2018YFC0910400), China Postdoctoral Foundation Grant (2017□M623188) and the Fundamental Research Fund of Xi’an Jiaotong University (1191329155). The authors also would like to thank anonymous reviewers for their comments and suggestions to improve this manuscript.

## CONFLICT OF INTEREST

The authors declare no conflict of interest.

## Notes

### Competing Interest Statement

The authors have declared no competing interest.

