## Supplementary figures and images for "Structural variants contribute to pangenome evolution of a plant pathogenic fungus"

### Figure S1

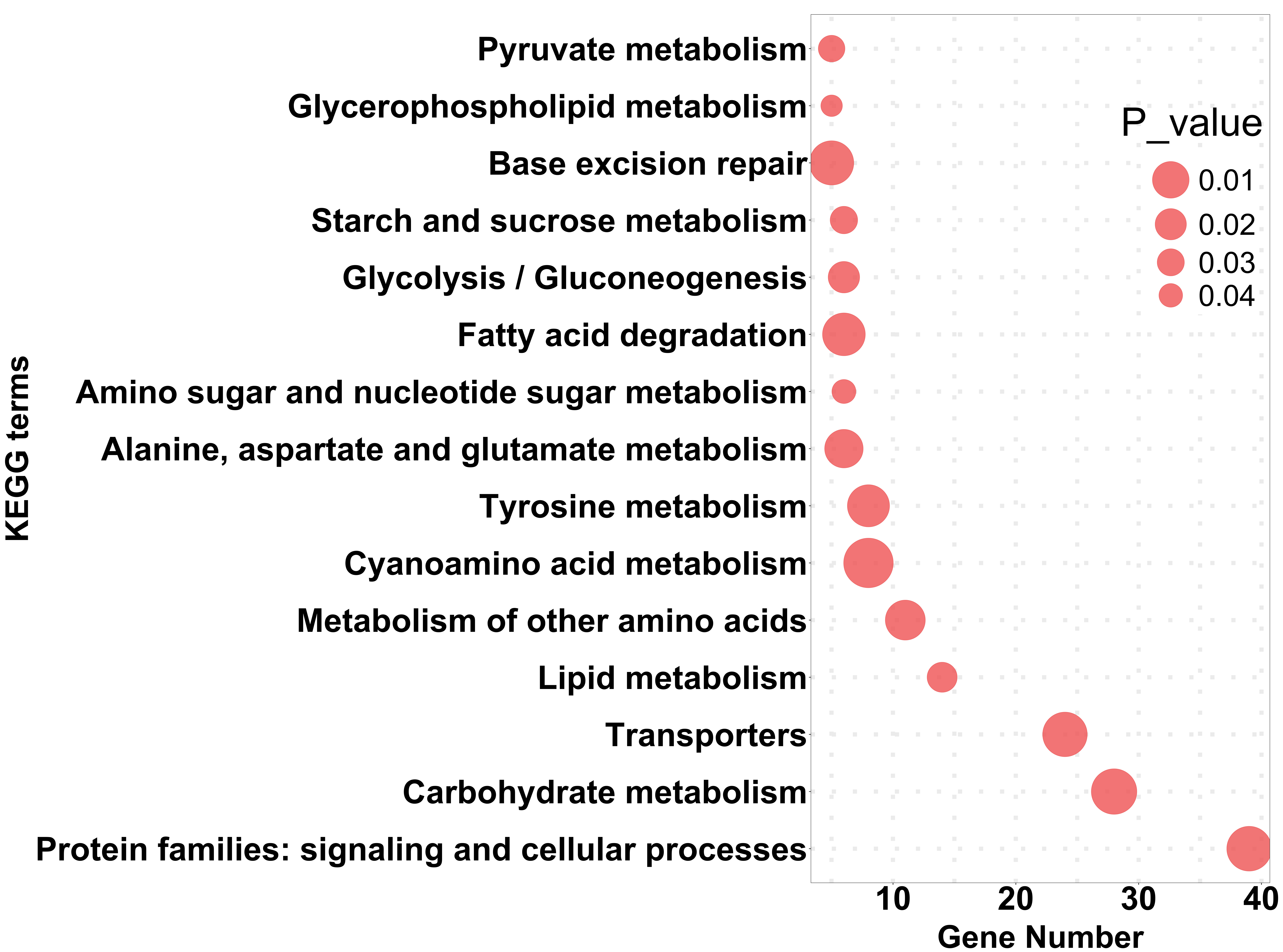

### Figure S2

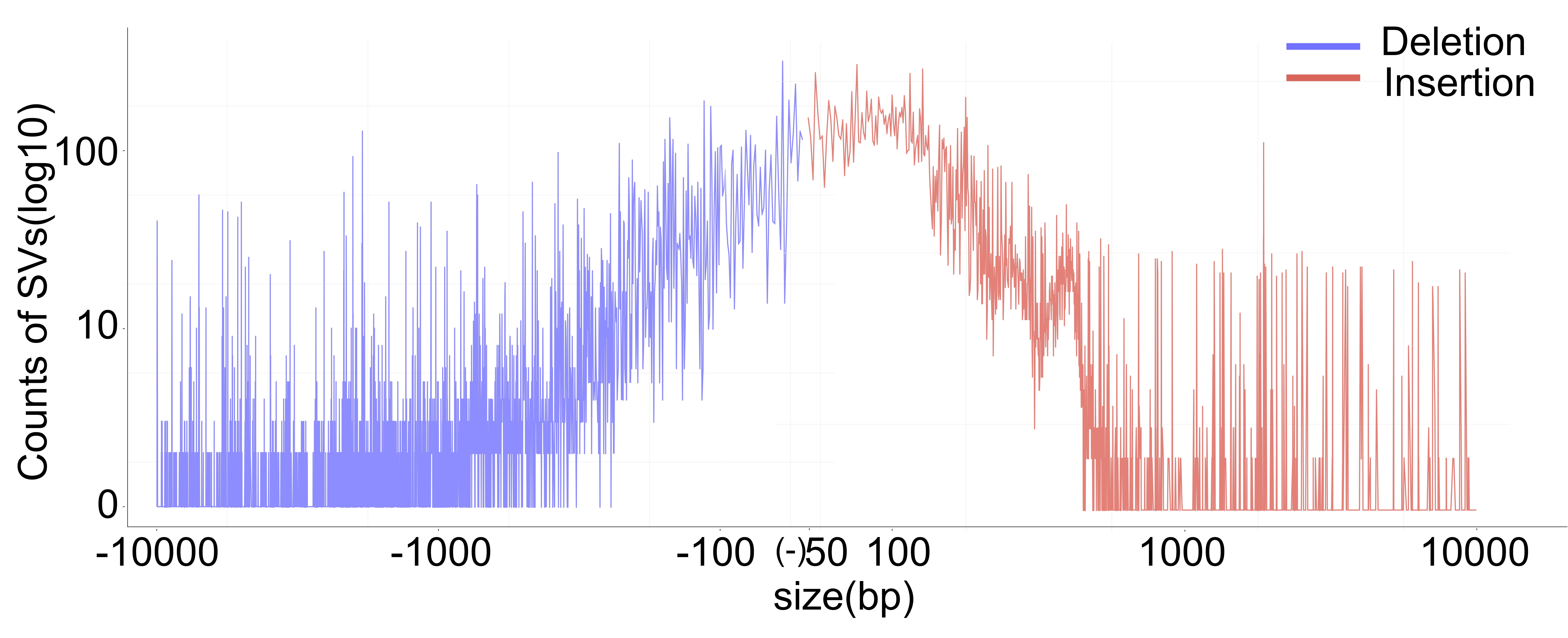

### Figure S3

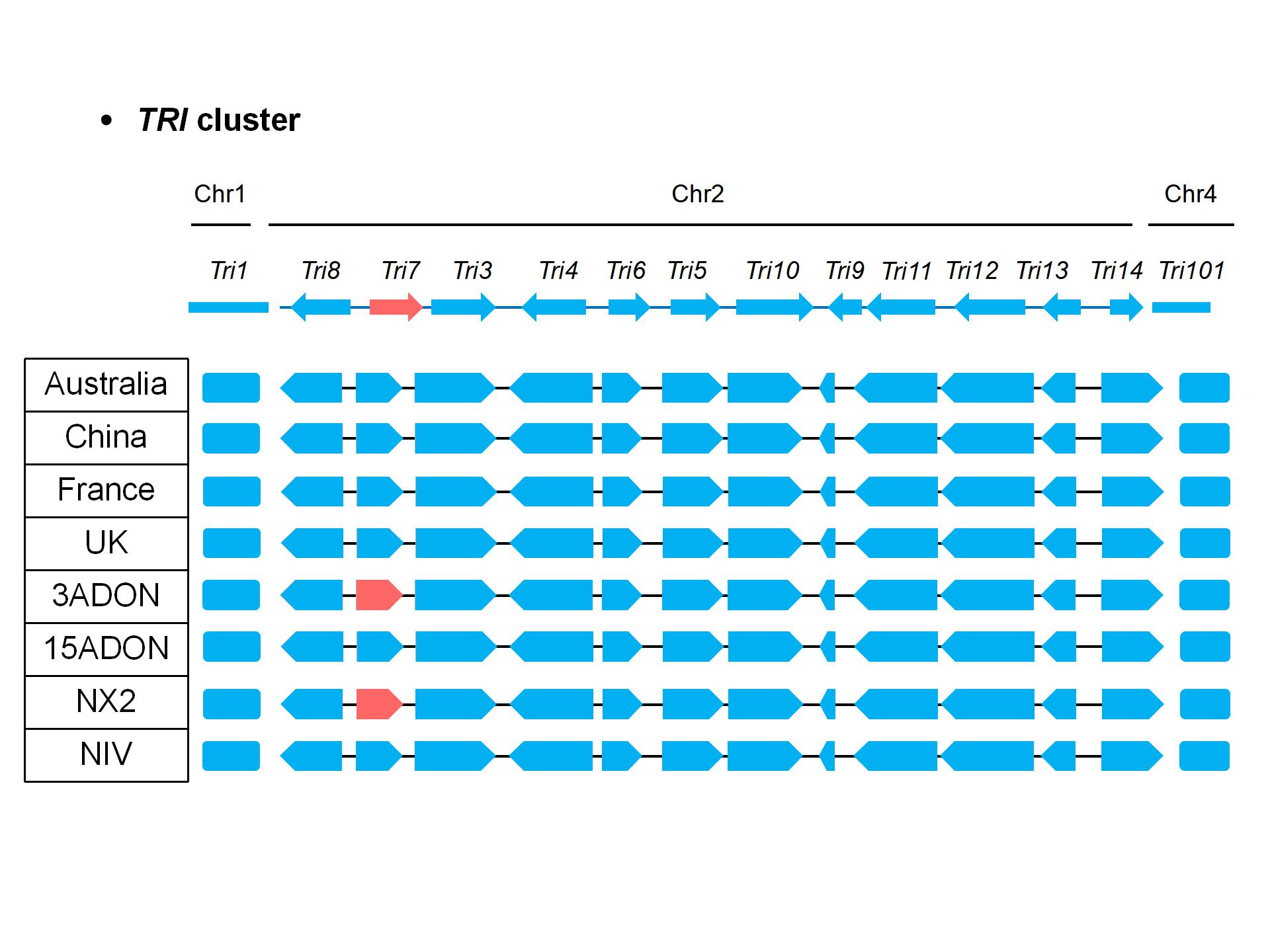

### Figure S4

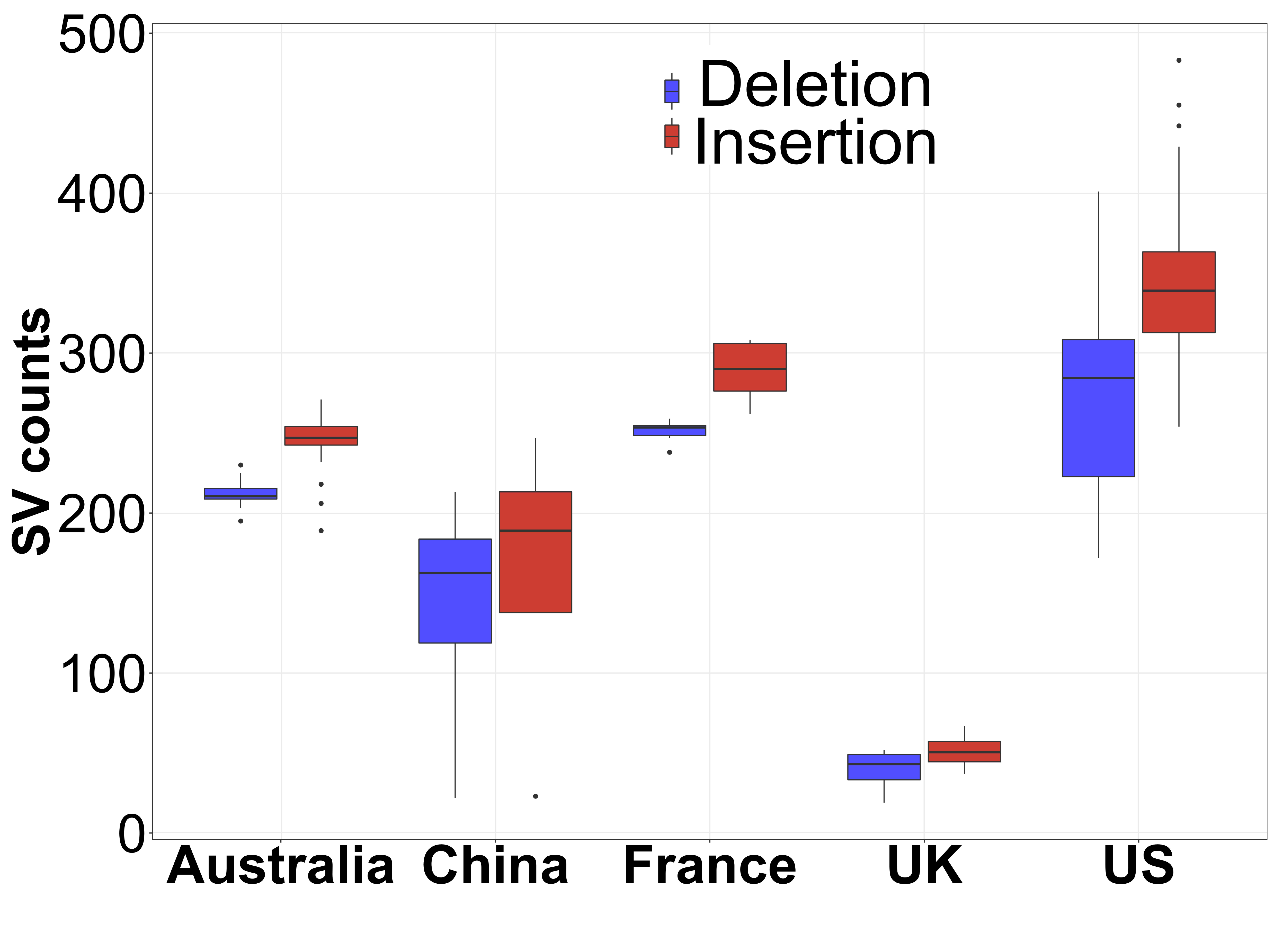

### Figure S5

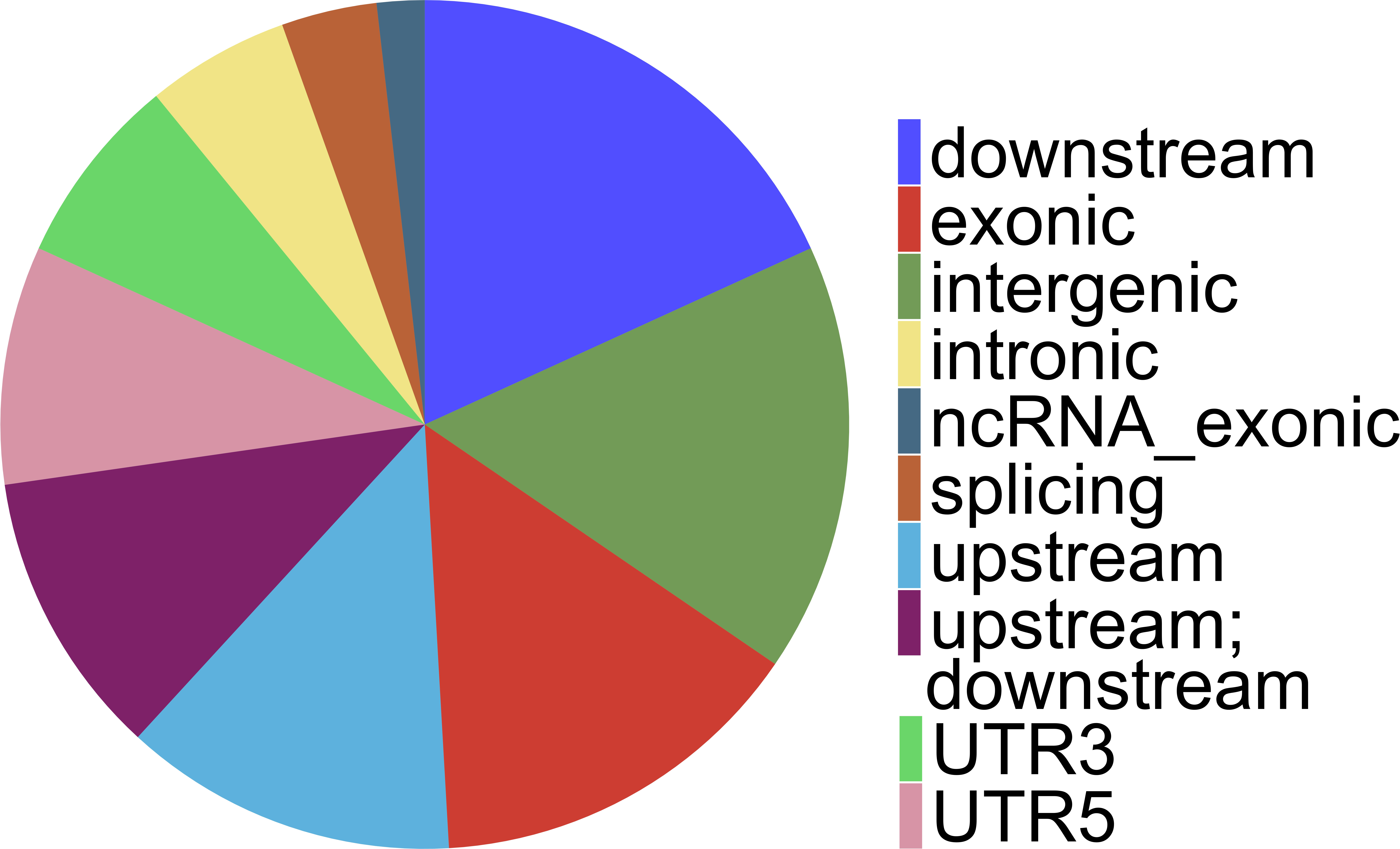
